# Cholangiocyte RUNX1 Orchestrates Fibrogenic and Inflammatory Signaling to Drive Biliary Fibrosis

**DOI:** 10.64898/2026.05.20.726667

**Authors:** Sayed Obaidullah Aseem, Jing Wang, Ahmed Younis, Diana Nakib, Grayson Way, Christiane Carter, Derrick Zhao, Yun-Ling Tai, Xuan Wang, Emily Gurley, Sonya MacParland, Phillip B. Hylemon, Nidhi Jalan-Sakrikar, Robert C. Huebert, Saul J. Karpen, Arun J. Sanyal, Huiping Zhou

## Abstract

**Introduction:** Biliary fibrosis and inflammation are central to the pathogenesis of cholangiopathies such as primary sclerosing cholangitis (PSC) and primary biliary cholangitis (PBC). Inflammatory and fibrogenic stimuli, such as transforming growth factor-β (TGFβ) and lipopolysaccharide (LPS) signaling, drive these processes, but their underlying transcriptional mechanisms in cholangiocytes remain incompletely defined. We investigated the role of Runt-related transcription factor 1 (RUNX1) as a transcriptional co-regulator of fibroinflammatory signaling in cholangiocytes.

**Methods:** Human PSC-derived cholangiocytes (PSC-Cs) and mouse large biliary epithelial cells (MLEs) were subjected to RUNX1 knockdown or pharmacologic inhibition (Ro5-3335 or AI-10-104). Cytokine secretion was profiled by Luminex multiplexing; RUNX1 genomic binding and protein interactome were assessed by ChIP-qPCR, ChIP-seq, and LC-MS/MS. *In vivo*, Mdr2^−/−^ mice received Ro5-3335, and cholangiocyte-selective Runx1 knockout mice (Krt19-CreERT) were challenged with a DDC diet, followed by evaluation of fibrosis and inflammation.

**Results:** RUNX1 expression was significantly increased in cholangiocytes from PSC and PBC patients, and Mdr2^−/−^ mice. RUNX1 knockdown or inhibition reduced IL6, TNFα, and other proinflammatory cytokines in PSC-Cs and attenuated TGFβ-, LPS-, and TNFα-induced Il6 and Ccl2 expression in MLEs. ChIP-qPCR and ChIP-seq revealed TGFβ-induced RUNX1 binding to the *Il6* promoter and 727 additional genomic sites enriched for fibrosis and inflammatory pathways; predicted upstream regulators included TGFβ, TNF, and NFκB signaling. Proteomic analysis identified TGFβ-induced RUNX1 interactions with SMAD2 and NFκB2. *In vivo*, Ro5-3335 treatment in Mdr2^−/−^ mice reduced hepatic collagen, ECM gene expression, immune cell infiltration, and serum liver injury markers and bile acids. Similarly, cholangiocyte-specific *Runx1* deletion mitigated fibrosis, inflammation, and liver injury in DDC-fed mice.

**Conclusion:** RUNX1 is a central transcriptional hub integrating TGFβ and inflammatory signals in cholangiocytes. Its inhibition attenuates biliary fibrosis and inflammation in cholestatic models, supporting RUNX1 as a potential therapeutic target in fibroinflammatory cholangiopathies.

## INTRODUCTION

Biliary fibrosis is the predominant pathological process in cholestatic liver diseases such as primary sclerosing cholangitis (PSC) and primary biliary cholangitis (PBC). These diseases collectively account for at least 16% of liver transplantation performed in the U.S. with an annual cost of $400 million [1]. They are also associated with considerable risk of hepatobiliary cancers. Morbidity and mortality closely correlate with biliary fibrosis. Despite this burden, there are currently no FDA-approved pharmacotherapies that halt or reverse biliary fibrosis, and late-stage cholangiopathies typically necessitate liver transplantation.

The pathophysiology of these diseases is driven by “reactive cholangiocytes” [2]. Following biliary injury, cholangiocytes transition from a quiescent epithelial state to a highly secretory, senescence-associated phenotype. These activated cholangiocytes secrete cytokines, chemokines, and growth factors that recruit immune cells and activate hepatic stellate cells (HSCs) and portal fibroblasts into collagen-producing myofibroblasts [2]. However, the molecular mechanisms of activated cholangiocytes that allow for its secretory phenotype and paracrine interactions are not completely understood.

A key pathway of fibrosis is transforming growth factor beta (TGFβ) signaling, a potent activator of hepatic stellate cells and portal fibroblasts into a myofibroblast-like phenotype that secretes extracellular matrix (ECM) [3]. Immune cells such as macrophages secrete TGFβ, which also activates cholangiocytes [4, 5]. Cholangiocytes activated by TGFβ recruit transcriptomic regulators, including epigenetic enzymes, which result in the expression of fibroinflammatory signals [6-8]. Activated cholangiocytes themselves secrete TGFβ as well as interleukin (IL)-6 and monocyte chemoattractant protein-1 (MCP1/CCL2), among other cytokines, further propagating the fibroinflammatory signaling [9] [10]. These signals recruit, differentiate and/or activate lymphocytes and myeloid cells, such as the autoimmune disease-associated T helper (Th) 17 cells and macrophages [11-13].

Our previous transcriptomic and epigenomic interrogation of TGFβ signaling revealed Runt-related transcription factors (RUNX) as a candidate downstream mediator of signaling in activated cholangiocytes [6]. These proteins are master regulators of embryonic development and are involved in the regulation of proliferation, differentiation, cell death and immune response [14, 15], interacting with TGFβ signaling pathway, among others [16, 17]. Mammals have three RUNX genes (RUNX1-3) with specific and dynamic expression patterns. RUNX1 has been ascribed conflicting roles in hepatobiliary fibrosis, both fibrogenic in hepatic stellate cells [18, 19] and antifibrogenic in hepatocytes under cholestatic conditions [20]. The role of RUNX transcription factors in cholangiocyte biology remains unexplored. Similarly, the involvement of RUNX1 in the production of fibroinflammatory signals through the TGFβ or inflammatory pathways is not well understood.

In this study, we identified a novel transcriptomic mechanism wherein RUNX1 acts as a central node in cholangiocyte activation, filling important gaps in the understanding of biliary fibrosis. RUNX1 directly regulated gene expression of inflammatory signals, such as IL6, downstream of TGFβ and inflammatory stimuli, both in human and mouse cholangiocytes. By utilizing pharmacological inhibition and cholangiocyte-specific genetic knockdown models, we demonstrate that RUNX1, upregulated in human cholestatic liver diseases, is a critical and druggable driver of biliary fibrosis and inflammation.

## MATERIALS AND METHODS

### Cell culture

Mouse large biliary epithelial cells (MLEs) were cultured in complete DMEM media [21, 22]. PSC patient-derived cholangiocytes (PSC-Cs) were kindly provided by Dr. Nicholas LaRusso (Mayo Clinic, Rochester, MN, USA) and were cultured in DMEM/F12 supplemented with 10% fetal bovine serum, 1% penicillin-streptomycin, adenine, insulin, epinephrine, T3-T, hydrocortisone and epidermal growth factor [23, 24]. Cells were serum starved before treatment with 10 ng/mL recombinant TGFβ (R&D Systems #240-B) for 16 hours. For siRNA targeting, cells were transfected with RUNX1 or control siRNAs (Dharmacon, ON-TARGETplus SMARTPool) using the Oligofectamine reagent (Invitrogen #12–252-011). The RUNX inhibitors, Ro5-3335 (Med Chem Express, HY-108470/CS-0028862) and AI-10-104 (Aobious, AOB17076) were dissolved in DMSO and cells treated at the indicated doses. For some experiments, nuclear extracts were isolated as previously described [25].

### Quantitative RT-PCR

Total RNA was extracted from cells using the TRIzol RNA Isolation Reagent (ThermoFisher, 15596026). Reverse transcription was performed with 2μg RNA using oligo (dT) primer and SuperScript III. Real-time PCR was performed using Sybr Green Master Mix in the QuantStudio™ 3 Real-Time PCR System (Applied Biosystems™) using primer sequences shown in Supplemental Table 1.

### ELISA

Mouse and human IL6 and MCP1 ELISA kits (eBioscience, Human IL6 Cat# 88-7066-88, Mouse IL6 Cat# 88-7064-88, Mouse MCP1 Cat#88-7391-88) were used to quantify IL6 and MCP1 in the cell cultured media following manufacturer protocols.

### Immunoblotting

Cell protein isolated in RIPA buffer were subjected to BCA protein quantification, electrophoresed on 4–20% Tris-Glycine gels, and transferred onto nitrocellulose membranes (Scientific Laboratory Supplies) for blotting. The membrane was blocked, incubated with primary antibodies, rinsed with TBST, and incubated with fluorophore conjugated secondary antibodies and imaged as described previously [25].

### Histochemistry and Immunofluorescence

Picrosirius red staining was performed on de-paraffinized liver sections following manufacturer protocols (Abcam, ab150681). Immunofluorescence was performed as described previously [25].

### Chromatin Immunoprecipitation (ChIP)

ChIP was performed using the SimpleChIP® Enzymatic Chromatin IP Kit (Cell Signaling, 9003S) following the manufacturer’s protocol. 15μg of DNA from nuclear extracts were incubated with 3-4 μg of primary antibody (RUNX1, ThermoFisher, PA5-19638) or IgG. Walking primers for the mouse *Il6* promoter were designed for quantitative PCR (Supplemental Table 2).

### ChIP-seq

40-180ng of DNA from 3 independent ChIP experiments were used to synthesize libraries using the 2S Plus DNA library kit from IDT. Sequencing was performed with Nextseq 2000 XLEAP. Reads were trimmed by Trimmomatic using a sliding window of 4 bp and minimum 20 phred score [26]. Alignment was performed using bowtie2’s (version 2.5.2) end-to-end setting to the GRC mm39 reference [27]; filtered to remove alignments under a quality score of 30, and multi-mappers. Peaks were called using MACS2. Significantly gained/lost peaks were defined as regions that had false discovery rate (FDR) values of less than 0.05, which excluded all but one peak with <1.5-fold change. Motif analysis was performed on the intersection of the biological replicates using Homer’s findMotifsGenome [28]. Peaks were visualized using integrated genome viewer (IGV). Genes associated with differential peak binding were investigated with downstream enrichment analysis using Qiagen Ingenuity Pathway Analysis (IPA). Library synthesis and sequencing was performed by the VCU Genomics Core. Bioinformatic analysis was performed by the VCU Massey Cancer Center Bioinformatics shared resource. The ChIP-seq raw data and meta data were deposited into the GEO Database (GSE306786).

### Immunoprecipitation (IP)

Nuclear protein co-immunoprecipitation was performed using the Nuclear Complex Co-IP Kit (Active Motif, 54001) using 3 μg RUNX1 antibody (ThermoFisher, PA5-19638) or IgG using 150-200 μg of nuclear protein.

### IP-Proteomic analysis

Proteomic analysis of RUNX1 nuclear immunoprecipitant was performed by liquid chromatography (LC)-MS/MS. IP proteins were first cleaned up and digested using the PreOmics iST kit following manufacturer protocol. The eluent was then placed in a vacuum evaporator at 45 deg C until completely dried. The LC-MS system consisted of a Thermo Electron Q-Exactive HF-X mass spectrometer with an Easyspray Ion source connected to an Acclaim PepMap 75μm x 2cm nanoviper C18 3μm x 100Å pre-column in series with an Acclaim PepMap RSLC 75μm x 50cm C18 2μm bead size (Thermo Scientific). Peptides were injected onto the column above. Peptides were eluted from the column with an 80%ACN 0.1% formic acid gradient at a flow rate of 0.3μl/min over 1 hour. The nano-spray ion source was operated at 1.9 kV. The digests were analyzed using the rapid switching capability of the instrument thus acquiring a full scan mass spectrum to determine peptide molecular weights followed by product spectra (40 High Energy C-trap Dissociation HCD spectra). This mode of analysis produces approximately 50,000 MS/MS spectra of ions ranging in abundance over several orders of magnitude. Not all MS/MS spectra are derived from peptides. The data were analyzed by database searching using the Sequest HT search algorithm with a custom mouse database downloaded from Swiss Prot. The data was processed with Proteome Discoverer 3.0.

### Cytokine array

Cultured media from PSC-Cs treated with control or RUNX1 specific siRNA were used for Luminex xMAP technology based multiplexed quantification of 96 Human cytokines, chemokines, and growth factors by Eve Technologies Corp (Calgary, Alberta).

### Mouse studies

Mdr2^−/−^ mice on C57BL/6J background were kindly provided by Dr. Daniel Goldenberg (Hadassah-Hebrew University Medical Center, Jerusalem, Isreal) [29]. 18-week-old Mdr2^−/−^ mice were administered Ro5-3335 at 50 mg/kg or vehicle only intraperitoneally every other day for four weeks. Separately, Mdr2^−/−^ on the FVB/N background were also similarly treated.

Runx1 floxed (Runx1^fl/fl^) mice were kindly provided by Dr. Nancy A. Speck (University of Pennsylvania, Philadelphia, PA, USA), which were crossed with mice containing tamoxifen-inducible Cre recombinase, driven by a cholangiocyte-specific promoter, Krt19-CreERT. Runx1^fl/fl^-Krt19-CreERT mice were intraperitoneally injected with tamoxifen at 80 mg/kg every other day for a week resulting in genetic knockdown of Runx1 in cholangiocytes (Runx1 CKO). As controls, Runx1^fl/fl^ without Krt19-CreERT were also similarly injected with tamoxifen resulting in wild-type expression of Runx1 (Ctrl). Mice were then subjected to a diet containing 0.1% 3,5-diethoxycarbonyl-1,4-dihydrocollidine (DDC) to induce cholangitis and biliary fibrosis with breaks of 2 nights of regular diet every week for 3 weeks. All mice continued to receive tamoxifen administration twice a week. At the end of the experiment, mice were euthanized, liver and blood harvested.

All mice were housed under 12-12-hour light and dark cycles with water and standard chow (or DDC diet) *ad libitum*. All mouse experiments were conducted in strict accordance with protocols approved by the Virginia Commonwealth University Institutional Animal Care and Use Committee.

### Human transcriptomic studies

Transcriptomic analysis was performed on previously published human PSC and PBC liver datasets available through the PSC sc/snRNAseq atlases and GEO database GSE243977 [30]. Data were processed and normalized using previously described pipelines [30]. Cholangiocyte populations were identified using canonical markers, and RUNX1 expression determined for each cell.

### Statistical analysis

Data are presented as mean with standard deviation. In mouse experiments, individual data points are also included in the graphs as dot plots. Student’s t-test was utilized to assess significance between two groups. Analysis of variance (ANOVA) with Tukey post-test or non-parametric Kruskal Wallis with Dunn’s post-test depending on the normality of the data was used to assess the statistical significance between more than 2 groups.

## RESULTS

### RUNX1 is overexpressed in cholangiopathies and essential for cholangiocyte expression of fibroinflammatory signals

Activated cholangiocytes are central to the pathogenesis of biliary fibrosis by virtue of a highly secretory phenotype mediating interactions with immune and ECM-producing myofibroblast cells. We have previously shown that cholangiocytes treated with TGFβ have increased expression of fibroinflammatory signals that are essential in activating myofibroblasts to produce the ECM of biliary fibrosis. With our previous transcriptomic and epigenomic interrogation of TGFβ signaling revealing RUNX transcription factors as candidate downstream mediators of signaling in activated cholangiocytes [6], we sought to assess their expression in PSC liver and mouse models. RUNX1 expression is significantly increased in the cholangiocytes from PSC and PBC livers as well as in the whole liver samples from PSC subjects in two separate databases (PSC/PBC sc/snRNAseq atlases and GEO database GSE243977 [30] and GSE159676) (Fig. 1A and Supplementary Table 1A/B). Similarly, mRNA expression of RUNX1 was significantly increased in cholangiocytes isolated from Mdr2^−/−^ (Fig. 1B).

**Fig 1.**
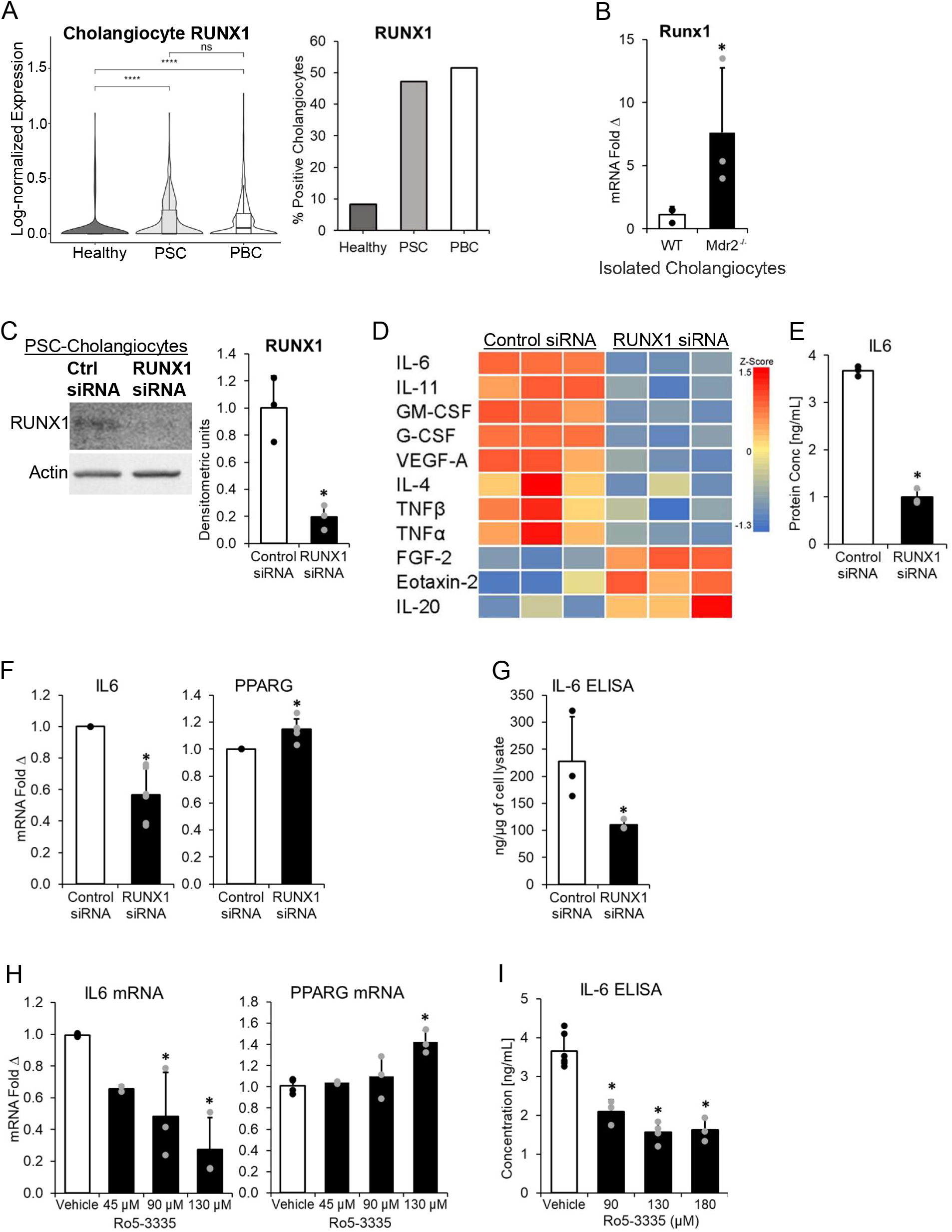
A) Transcriptomic analysis of PSC and PBC liver samples, compared with healthy livers, showed significantly increased RUNX1 expression in cholangiocytes, as well as an increased fraction of RUNX1-expressing cholangiocytes (PSC/PBC sc/snRNAseq atlases and GEO database GSE243977). B) qPCR analysis of cholangiocytes from Mdr2^−/−^ mice compared to wild type (WT). C) In PSC-Cs, RUNX1 expression was significantly knocked down with specific siRNAs as demonstrated by Western blot analysis. D) Heatmap of significantly altered levels of cytokines in the media of PSC-Cs with RUNX1 knockdown using Luminex xMAP technology based multiplexed quantification. E) IL6 levels from D. F) qPCR analysis of *IL6* and *PPARG* expression with RUNX1 knockdown. G) ELISA assay demonstrating levels of IL6 in the media of control and RUNX1 siRNA treated PSC-Cs. H) qPCR analysis of *IL6* and *PPARG* levels showed dose dependent response to RUNX1 inhibitor, Ro5-3335. I) IL6 ELISA of media treated with Ro5-3335. (N=3 at least, *= p<0.05)

Next, we utilized RUNX1 specific siRNA to significantly knockdown its expression by >80% (Fig. 1C) in cholangiocytes obtained from PSC patients (PSC-Cs). The effect of RUNX1 knockdown on cytokines secreted into the cultured media was assessed using a commercial 96 cytokine immunoarray (Luminex xMAP technology, by Eve Technologies Corporation) (Supplementary Table 2). Several inflammatory markers were significantly attenuated with RUNX1 knockdown including IL6 and its family member IL11 along with tumor necrosis factors (TNF) as well as IL4 and granulocyte-macrophage colony-stimulating factor (Fig. 1D and E). RUNX1 knockdown resulted in significant reductions in IL6 mRNA and protein secretion by ELISA of cultured media (Fig. 1F and G). Conversely, the expression of anti-fibroinflammatory factor, peroxisome proliferator-activated receptor gamma (*PPARG*), was increased (Fig. 1F). Similarly, treatment of PSC-Cs with a RUNX1 inhibitor (Ro5-3335) resulted in significant attenuation of IL6 mRNA and protein in a dose dependent manner, while *PPARG* expression was increased (Fig. 1H and I).

### RUNX1 inhibition attenuates the TGFβ-stimulated fibroinflammatory signals

To determine the role of RUNX1 in the TGFβ-induced production of fibroinflammatory signals by cholangiocytes, we used the RUNX1 inhibitors, Ro5-3335 and AI-10-104. Ro5-3335 in mouse large biliary epithelial cells (MLEs) led to a dose-dependent attenuation of TGFβ-induced *Il6* and Monocyte chemoattractant protein-1 (*Mcp1/Ccl2*) mRNA (Fig. 2A). Similar findings were observed at the protein level in the cultured media using specific ELISAs for IL6 and MCP1 (Fig. 2B). Conversely, the expression of *Pparg*, was increased with the RUNX1 inhibitor treatment (Fig. 2A). Similar, dose-dependent findings were noted with AI-10-104 (Supplementary Fig. 1A and B).

**Fig 2.**
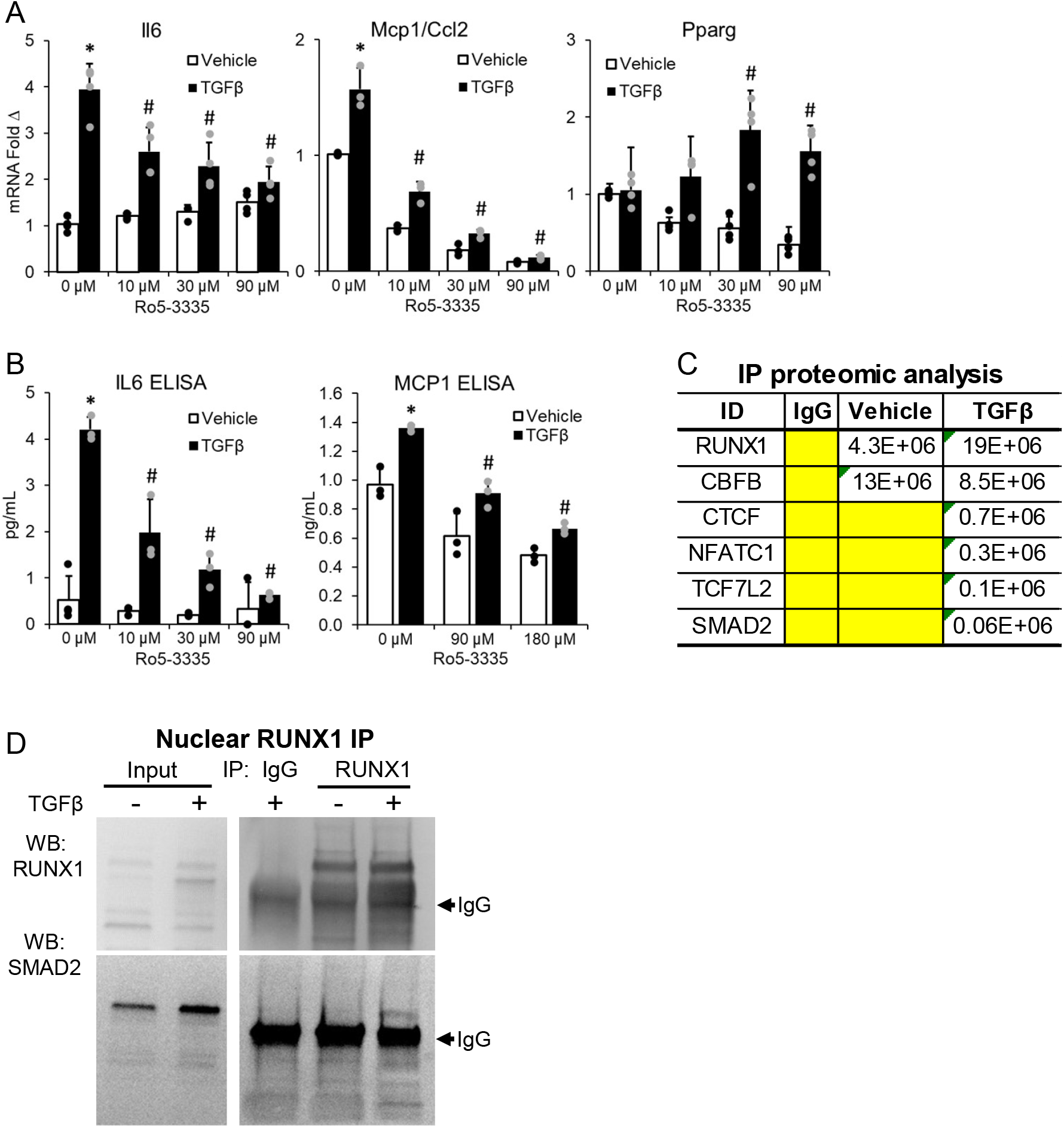
In MLEs, the TGFβ-induced elevated expression of *Il6* and *Ccl2/Mcp1* expression and protein level were dose-dependently attenuated with the RUNX1 inhibitor, Ro5-3335, co-treatment (A-B). C) Proteomic analysis of RUNX1 IP by mass spectrometry demonstrated interactions with SMAD2 and additional transcriptional co-regulators. Protein abundance was semi-quantitatively estimated based on the number of unique peptides identified. D) RUNX1 specific IP showed direct nuclear interaction with SMAD2 following TGFβ stimulation. (N=3 at least, # p<0.05 when compared to TGFβ only)

To determine whether RUNX1 interacted with the TGFβ canonical SMAD signaling and to identify co-transcription factors, we employed immunoprecipitation (IP) following TGFβ stimulation. RUNX1 IP was subjected to unbiased proteomic analysis via liquid chromatography with tandem mass spectrometry (LC-MS/MS), followed by semi-quantitative estimation of protein abundance based on the number of unique peptides (Fig. 2C). RUNX1 abundance was significantly enriched with TGFβ treatment. The RUNX binding partner, core binding factor beta (CBFB), was among the most enriched interacting co-factors, but its levels did not significantly change with TGFβ stimulation. Other prominent co-factors included SMAD2 and others (Fig. 2C). These transcription factors had significantly enriched binding sites in our previously performed SMAD3 ChIP-seq of TGFβ-stimulated cholangiocytes (Supplementary Fig. 1C) (GSE145253) [6]. Canonical regulators of inflammation (such as NFKB1 or STAT1/3) were not identified to significantly interact with RUNX1. However, NFKB2 was detectable in the RUNX1 immunoprecipitant, albeit at very low levels, suggestive of a weak or transient interaction (data not shown). To validate the RUNX1 interactions with TGFβ signaling, RUNX1 IP was analyzed by Western blot demonstrating association with SMAD2 in nuclear extracts of cells treated with TGFβ compared to vehicle (Fig. 2D). These findings indicate that RUNX1 is essential in mediating the downstream effects of TGFβ signaling, interacting with other transcriptional co-regulators in stimulating fibroinflammatory signals.

### RUNX1 binds the promoters of fibroinflammatory genes upon TGFβ stimulation of cholangiocytes

To determine direct regulation of *Il6* expression by RUNX1, we performed chromatin immunoprecipitation (ChIP) for RUNX1. RUNX1 binding of *Il6* promoter was significantly increased with TGFβ stimulation of MLEs (Fig. 3A). Walking qPCR primers along the *Il6* promoter showed RUNX1 binding to the promoter region 200bp and upstream which was markedly increased (~2.5-fold) with TGFβ stimulation (Fig. 3A & B). Furthermore, the TGFβ-stimulated increased RUNX1 binding to the *IL6* promoter was attenuated to basal levels with the RUNX1 inhibitor (Ro5-3335) (Fig. 3A & B). These results suggest that RUNX1 directly regulates the expression of *Il6* stimulated with TGFβ.

**Fig 3.**
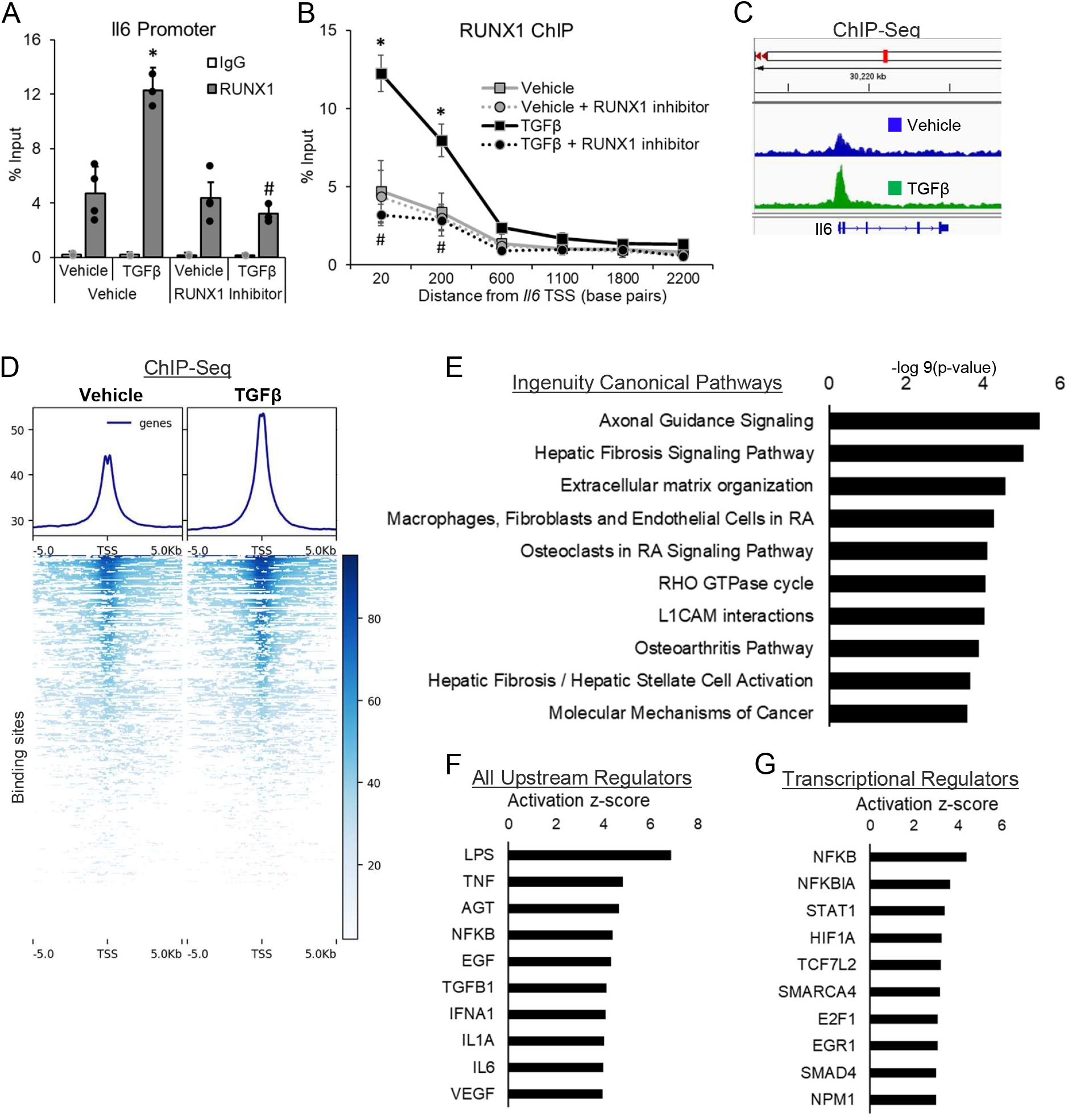
(A-B) ChIP-qPCR showed significantly increased RUNX1 binding to the proximal but not distal *Il6* promoter following TGFβ stimulation. The RUNX1 binding is significantly attenuated with the RUNX1 inhibitor, Ro5-3335. C-D) ChIP-seq showed increased TGFβ-induced RUNX1 binding of *ll6* and 727 other binding sites. E) Ingenuity Pathway Analysis of genes with increased TGFβ-induced RUNX1 binding showed enrichment in fibroinflammatory pathways. F-G) Putative upstream regulators of genes with increased RUNX1 binding included TGFβ and inflammatory mediators/pathways. (N=3 at least, # p<0.05 when compared to TGFβ only)

To evaluate the full spectrum of genes that might be regulated by RUNX1 binding of their promoters, we performed ChIP-seq analysis of MLEs stimulated with TGFβ overnight at 10 ng/mL and compared to vehicle treatment. Figure 3C shows the RUNX1 binding peaks localized at the promoter region of *Il6* in the ChIP-seq analysis. Upon TGFβ treatment, RUNX1 binding was significantly increased at 727 DNA sites compared to vehicle (Fig. 3D). The genes associated with sites that showed a statistically significant change in RUNX1 binding were identified. The Qiagen Ingenuity Pathway Analysis (IPA) was performed on this set of genes. The most significantly enriched pathways included matrix organization, hepatic fibrosis and other inflammatory pathways (Fig. 3E). Interestingly, while TGFβ and SMAD transcriptional regulators were among the most activated upstream regulators, the top candidates included upstream inflammatory and transcriptional regulators such as LPS, tumor necrosis factor (TNF) and NFKB (Fig. 3F and G). Taken together, these observations indicate RUNX1 regulating a set of fibroinflammatory genes within the TGFβ-stimulated genes.

### RUNX1 inhibition attenuates inflammatory stimuli-induced IL6 and MCP1 expression

Our above findings indicated that RUNX1 may serve as a key node in integrating inflammatory and fibrogenic signals. Since LC-MS/MS proteomic analysis of RUNX1 immunoprecipitate detected NFKB2 at low levels, we further investigated the possible interaction of RUNX1 with NFKB2. Indeed, RUNX1 IP of MLEs revealed the presence of NFKB2 following TGFβ stimulation (Fig. 4A). Furthermore, both TGFβ and TNFα stimulation resulted in significantly increased nuclear levels of NFKB2, indicating its activation by either stimulus (Fig. 4B). There was concomitant increased nuclear level of RUNX1 with TGFβ stimulation along with its canonical SMAD2 (Fig. 4B).

**Fig 4.**
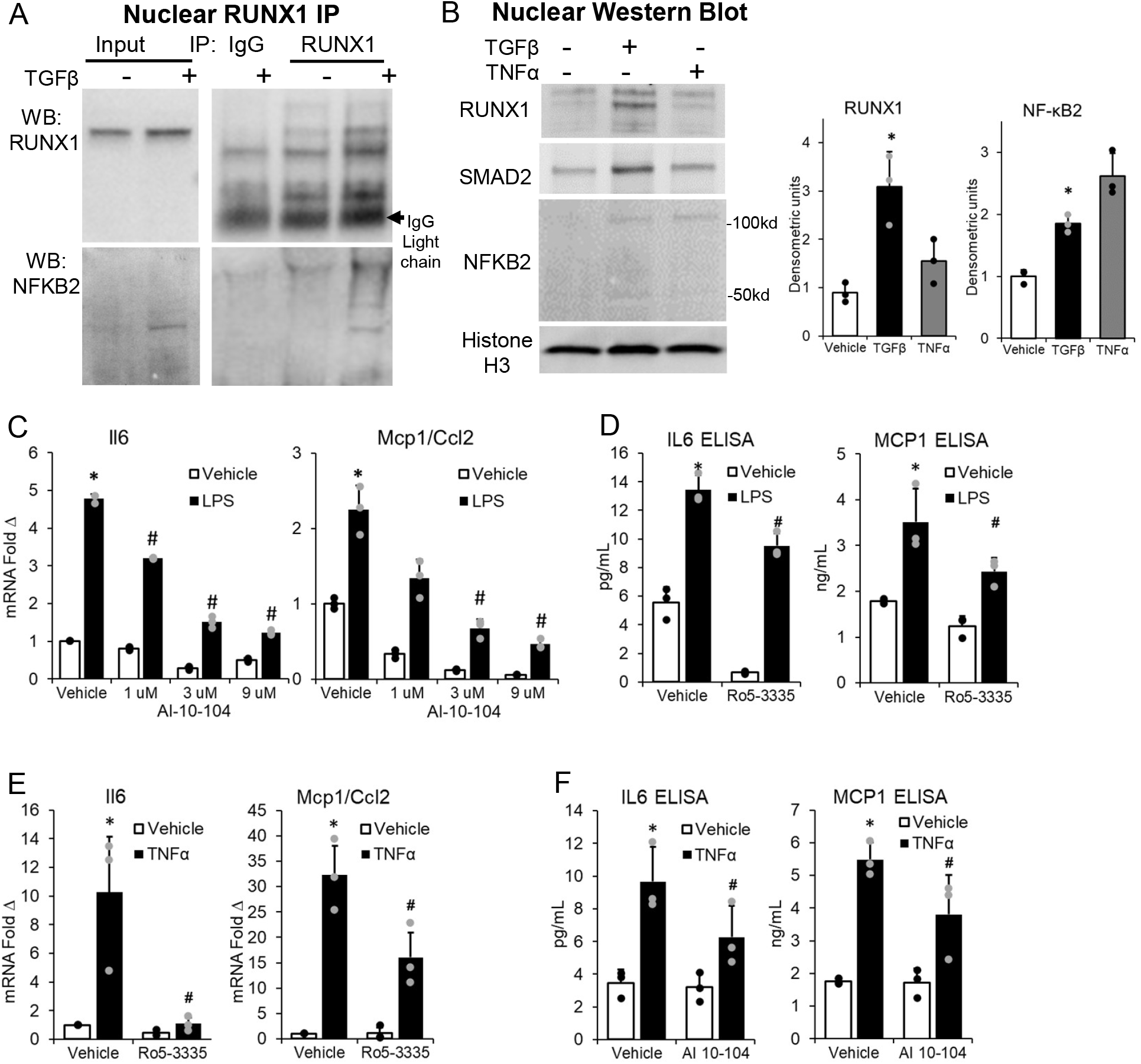
A) In MLEs, RUNX1 IP demonstrated co-IP of nuclear NFKB2 with TGFβ stimulation. B) Western blot analysis demonstrated increased nuclear NFKB2 p52 subunit and heterodimer with TGFβ or TNFα treatment. Nuclear RUNX1 was also significantly increased with TGFβ stimulation. C) The LPS-stimulated (100 ng/mL) expression of both *Il6* and *Ccl2* were significantly and dose-dependently attenuated by the RUNX1 inhibitor, AI-10-104. D) ELISA analysis demonstrated that the LPS-stimulated secretion of IL6 and MCP1 in the cultured media was significantly attenuated by the RUNX1 inhibitor, Ro5-3335 (180 µM). E and F) TNFα-stimulated expression of *Il6* and *Ccl2* were significantly attenuated at the mRNA and protein levels with the AI-10-104 inhibitor. (N=3 at least, # p<0.05 when compared to treatment group only)

To investigate the role of RUNX1 downstream of canonical activators of NFKB signaling, MLEs were treated with LPS and TNFα with and without RUNX1 inhibitors. The LPS-induced increase in the expression of IL6 and MCP1 were both significantly attenuated by RUNX1 inhibitors (both Ro5-3335 and AI-10-104) at the mRNA and protein levels (Fig. 4C and D). Combination of TNFα (at 5 and 10 ng/mL) and Ro5-3335 (at 90 and 180 μM) demonstrated significant cytotoxicity in MLEs. However, AI-10-104 was well tolerated and significantly attenuated the TNFα-stimulated increased expression of IL6 and MCP1 at both mRNA and protein levels (Fig. 4E and F). Together, these observations indicate that RUNX1 interacts with downstream regulators of both TGFβ and inflammatory stimuli and is essential in the expression of fibroinflammatory signals.

### RUNX1 inhibition in Mdr2^−/−^ mice results in reduced biliary fibrosis and markers of inflammation

Next, to determine if RUNX1 inhibition mediated reductions in cholangiocyte fibroinflammatory signals ameliorated peri-portal inflammation and fibrosis, 18-week-old Mdr2^−/−^ mice on the C57/B6 background were treated with the RUNX1 inhibitor (Ro5-3335) at 50 mg/kg by intraperitoneal injections every other day for 4 weeks. The mice treated with Ro5-3335 showed a trend toward more weight loss (Supplementary Fig. 2A). Histological analysis with picrosirius red staining showed significantly reduced staining with Ro5-3335 (Fig. 5A). Similarly, hepatic collagen quantified by hydroxyproline content by mass spectrometry demonstrated a significant reduction with Ro5-3335 treatment (Fig. 5B). Myofibroblast activation was significantly reduced with Ro5-3335 inhibition as demonstrated by immunofluorescence using αSMA marker (Fig. 5C). Similarly, macrophage peri-portal accumulation (determined by F4/80 staining) was significantly reduced with Ro5-3335 treatment (Fig. 5C). The area of CK7-positive bile ducts and surrounding infiltration, termed portal expansion, was significantly reduced with Ro5-3335 treatment, which is indicative of reduced ductular reaction (Fig. 5C). Hepatic mRNA expression of ECM components, smooth muscle actin (*Acta2*) and collagen (*Col1a1*), were significantly reduced with Ro5-3335 treatment (Fig. 5D). The abundance of the master regulator of inflammation, *Nfkb1*, was significantly reduced in the liver of mice treated with Ro5-3335 (Fig. 5E). The reduction in the expression of *Tnfa* with Ro5-3335 treatment approached statistical significance (Fig. 5E). The expression of *Il6* and *Ccl2* were generally low, varied widely and did not demonstrate a statistically significant difference, although the levels were generally lower with Ro5-3335 treatment (Fig. 5E). Likewise, we did not detect a change in liver injury markers, alkaline phosphatase (ALP) and alanine aminotransferase (ALT), which were only modestly elevated (Supplementary Fig. 2B).

**Fig 5.**
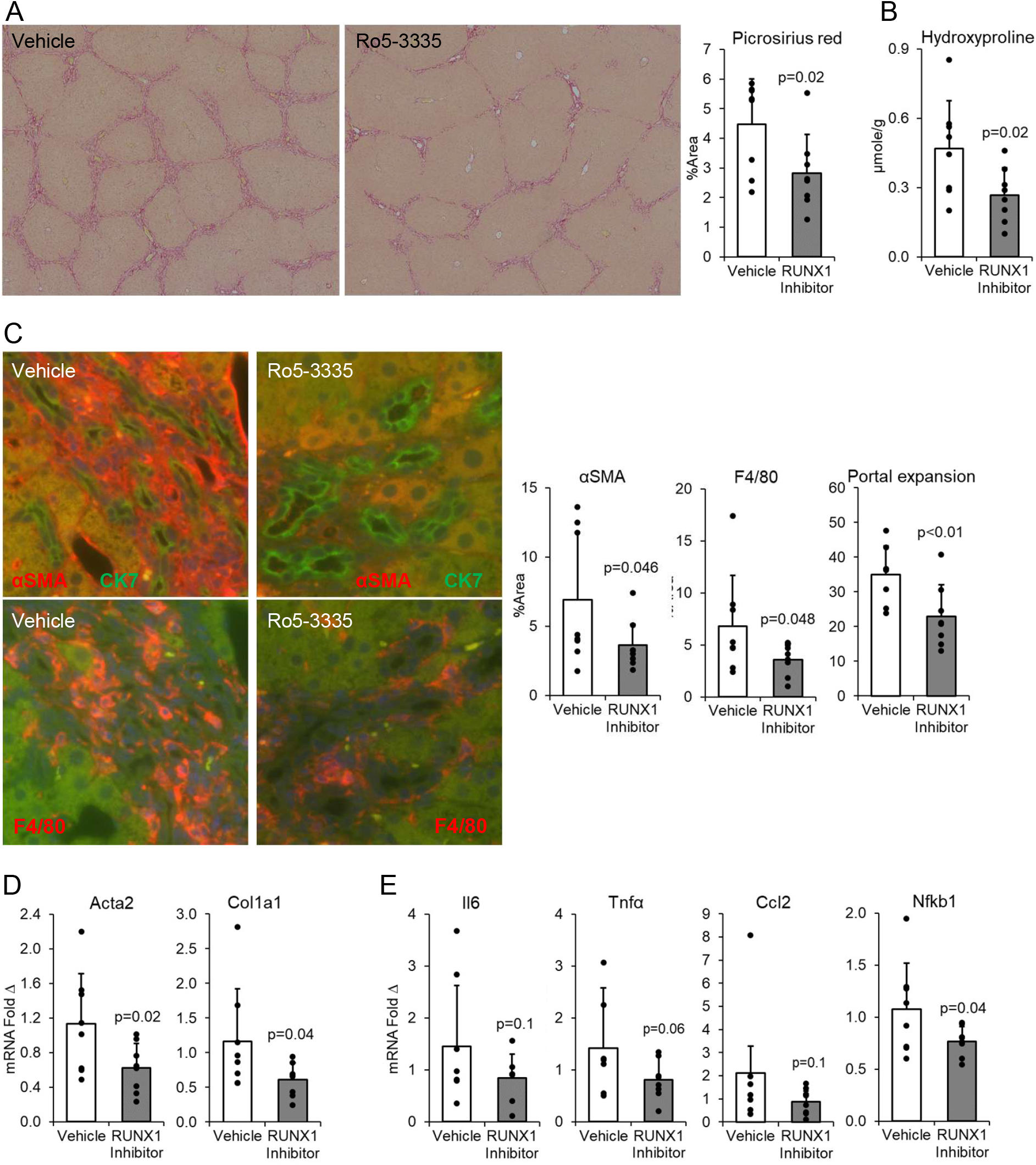
A) Picrosirius red staining of livers from Mdr2^−/−^ mice on the C57/B6 background showed reduced staining with Ro5-3335 treatment compared to vehicle with area quantification using ImageJ. B) Hydroxyproline content in the liver quantified by mass spectrometry was significantly reduced with Ro5-3335 treatment compared to vehicle. C) IF of αSMA and macrophage marker, F4/80, in sister sections showed reduced immunostaining surrounding CK19 immunostained bile ducts (green) along with reduced portal expansion with Ro5-3335 treatment. D&E) qPCR analysis of liver lysate demonstrated attenuation of ECM components, *aSMA/Acta2* and *Col1a1*, and inflammatory mediators, *Tnfa*, and *Nfkb*, with Ro5-3335 treatment. (N=8 per group).

Given the modest increase in inflammatory signals accompanied with only mild elevations in serum liver injury markers in the C57/B6 Mdr2^−/−^ mice, we aimed to further evaluate the *in vivo* role of RUNX1 in inflammation associated with biliary injury in Mdr2^−/−^ mice on the FVB/N background, known to develop more robust hepatic inflammation [31-33]. These mice were similarly treated with the Ro5-3335 inhibitor. Hepatic mRNA expression of inflammatory markers, *Il6, Ccl2, Tnfa* and *Nfkb* were all significantly reduced in mice treated with the RUNX1 inhibitor, but not the anti-inflammatory *Il10* (Fig. 6A). Histological analysis with hematoxylin and eosin (H&E) staining demonstrated markedly reduced accumulation of inflammatory cells surrounding the portal triads (Fig. 6B). Similarly, immunofluorescence analysis of liver sections using antibodies for immune cells, Cd45, and macrophages, F4/80, showed reduced peri-portal accumulation of immune cells in general and macrophages specifically (Fig. 6C). This reduction in inflammation was accompanied by improvement in cholestasis demonstrated by reduced serum total bile acid level and reduced hepatic injury markers, ALT and AST, but not ALP (Fig. 6D). The serum injury markers were significantly more elevated in this strain compared to those on the C57/B6 background (Supplementary Fig. 2B).

**Fig 6.**
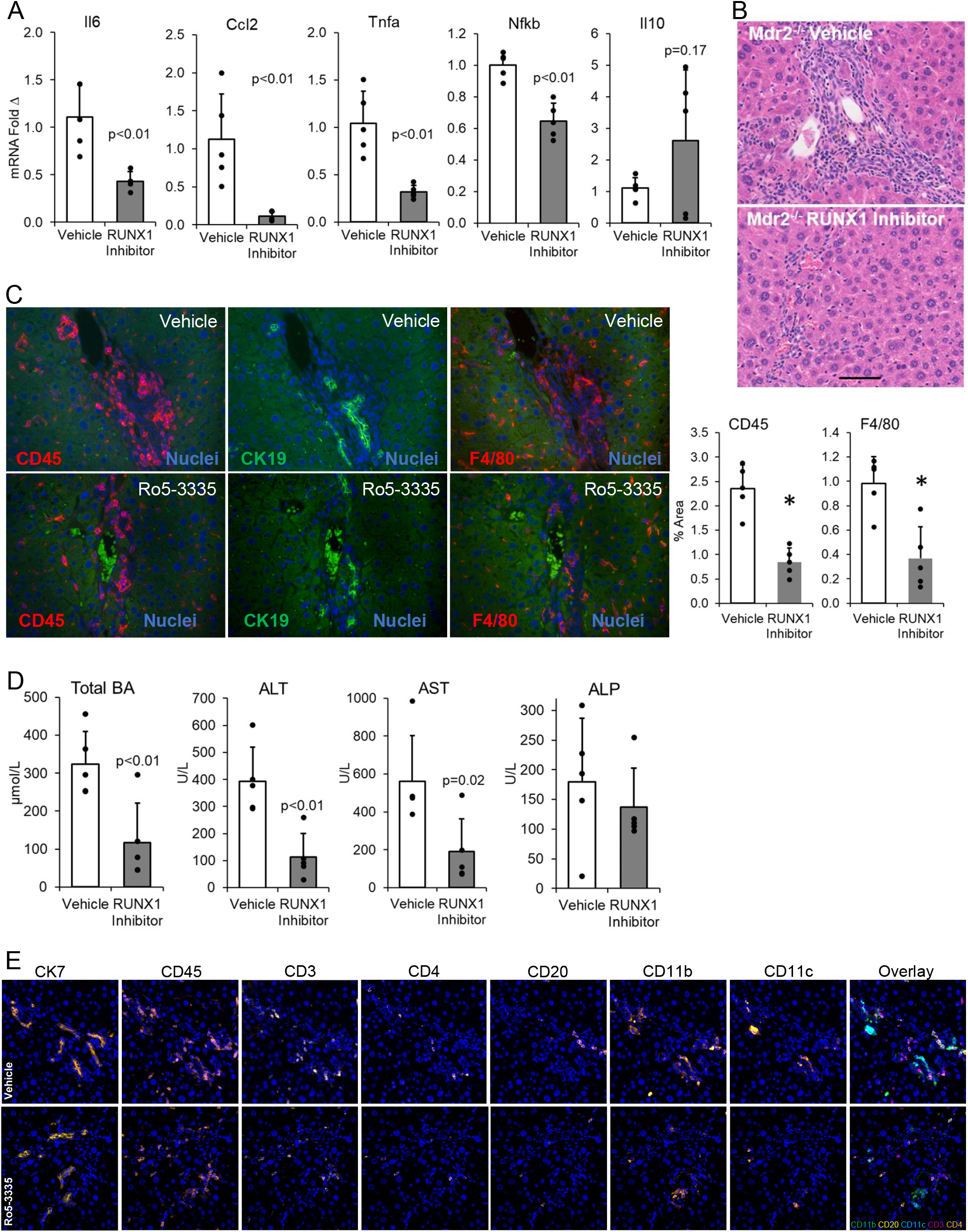
A) qPCR analysis of liver lysate from Mdr2^−/−^ mice on the FVB/N background treated with Ro5-3335 demonstrated attenuation of inflammatory mediators, *Il6, Ccl2, Tnfa* and *Nfkb*, but not the anti-inflammatory cytokine, *Il10* compared to vehicle. B) H&E stain showed reduced accumulation of immune cells in the hepatic peri-portal regions of Ro5-3335 treated mice. C) IF of a marker of immune cells, CD45, and macrophage marker, F4/80, in sister sections showed reduced immunostaining surrounding CK19 labeled bile ducts (green). D) Serum AST, ALT, and total bile acids were significantly attenuated with Ro5-3335 treatment. E) IF multiplexing of cholangiocyte marker (CK7) and immune cells demonstrated reduced periportal CD4+, CD20+, and CD11c+ cells with Ro5-3335 treatment but not CD3+ and CD11b cells. (N=5 per group).

To further delineate the types of immune cells and their periportal accumulation, we performed immunofluorescence-based multiplexing for immune markers to identify immune cell types in vehicle and Ro5-3335 treated mouse livers. While T cells (CD3+, CD45+) collectively were not significantly different, CD4+ T cells were significantly reduced in the liver of Ro5-3335 treated mice (Fig. 6E and Supplementary Fig. 2C). Similarly, B cells (CD20+) and ITGAX (CD11c+) myeloid cells were significantly reduced with Ro5-3335 treatment (Fig. 6E and Supplementary Fig. 2C). In contrast, IGAM (CD11b+) myeloid cells were not significantly different in the treatment compared to control group. Therefore, the reduction in periportal immune cells with Ro5-3335 was driven by reductions in CD4+ T cells, B cells and ITGAX myeloid cells. Taken together, these observations indicate that inhibiting RUNX1 results in significant reductions in biliary inflammation and fibrosis in the Mdr2^−/−^ mice.

### Cholangiocyte-specific deletion of *Runx1* results in reduced biliary fibrosis and markers of inflammation in the DDC diet mouse model

Given the protective effects of RUNX1 inhibitor, we sought to further investigate the role of *Runx1* in cholangiocytes *in vivo*. Thus, we generated a cholangiocyte-selective *Runx1* knockdown model by crossing *Runx1* floxed mice with a tamoxifen-inducible *Krt19-Cre* mice. Following tamoxifen treatment, *Runx1*^*fl/fl*^*;Krt19-Cre*^*+*^(*Runx1* KO) and *Runx1*^*fl*/fl^ (control) mice were subjected to DDC diet induced biliary injury, inflammation and fibrosis. qPCR analysis demonstrated increased expression of *Runx1* in the whole liver of mice on the DDC diet, which was significantly reduced in the mice with cholangiocyte-specific *Runx1* KO (Fig. 7A). *Acta2*, a marker of activated myofibroblasts, which was significantly increased with DDC diet, was attenuated with *Runx1* KO (Fig. 7A). Hepatic collagen content assayed by hydroxyproline quantification by mass spectrometry was significantly attenuated in *Runx1* KO mice on the DDC diet compared to controls (Fig. 7B). Similarly, picrosirius red staining of liver sections, which demonstrated increased staining with DDC diet, was significantly attenuated in *Runx1* KO mice on DDC diet (Fig. 7C). Furthermore, immunofluorescence analysis demonstrated significant attenuation of periportal and peri-ductular αSMA staining in *Runx1* KO mice on DDC diet compared to controls (Fig. 7D). The liver injury marker in serum, ALT, was significantly attenuated in *Runx1* KO mice on DDC diet compared to controls (Fig. 7E). Consistently, hepatic mRNA expression of inflammatory mediators (*Ccl2* and *Nfkb*) were significantly reduced in *Runx1* KO mice on DDC compared to controls, while *Il6* levels approached a statically significant reduction in *Runx1* KO mouse livers. Overall, these observations indicate that cholangiocyte-specific *Runx1* KO results in significant reductions in biliary fibrosis and markers of inflammation.

**Fig 7.**
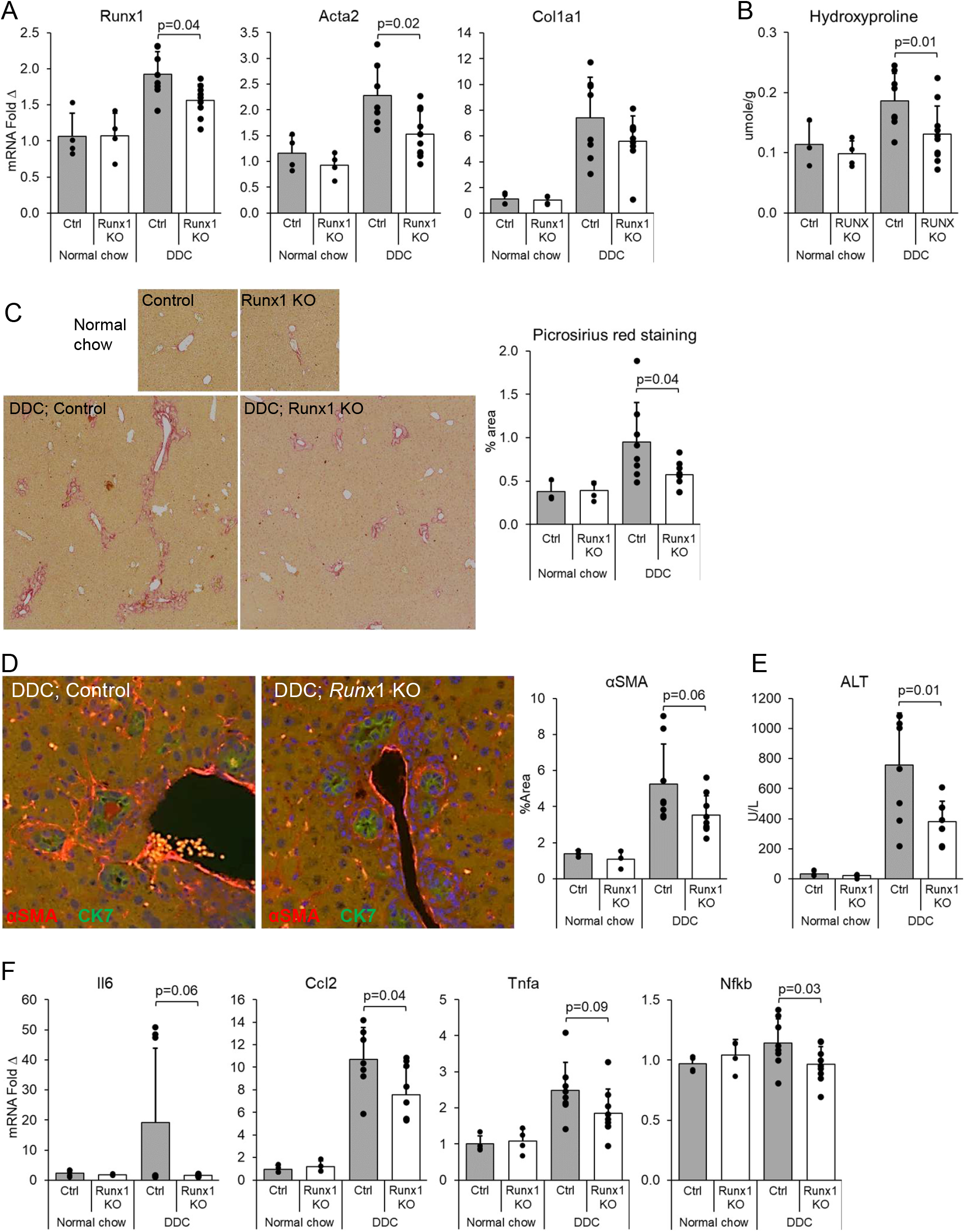
A) qPCR analysis of whole liver lysate of mice with cholangiocyte-specific *Runx1* deletion (*Runx1* KO) demonstrated reduced *Runx1* and *Acta2* expression compared to control mice following DDC diet. B) Hydroxyproline content in the liver measured by mass spectrometry was significantly reduced in *Runx1* KO compared to control mice on DDC diet. C) Picrosirius red staining of livers from *Runx1* KO mice showed reduced staining compared to control on DDC diet with area quantification using ImageJ. D) IF of αSMA appeared to show reduced immunostaining surrounding CK19 labeled bile ducts (green), approaching statistical significance. E) Serum ALT, a liver injury marker, was significantly reduced in *Runx1* KO compared to control mice. F) qPCR analysis of liver lysate demonstrated significant attenuation of *Ccl2* and *Nfkb* in *Runx1* KO. (N=4 per regular chow group, 8 per DDC diet group).

**Fig 8.**
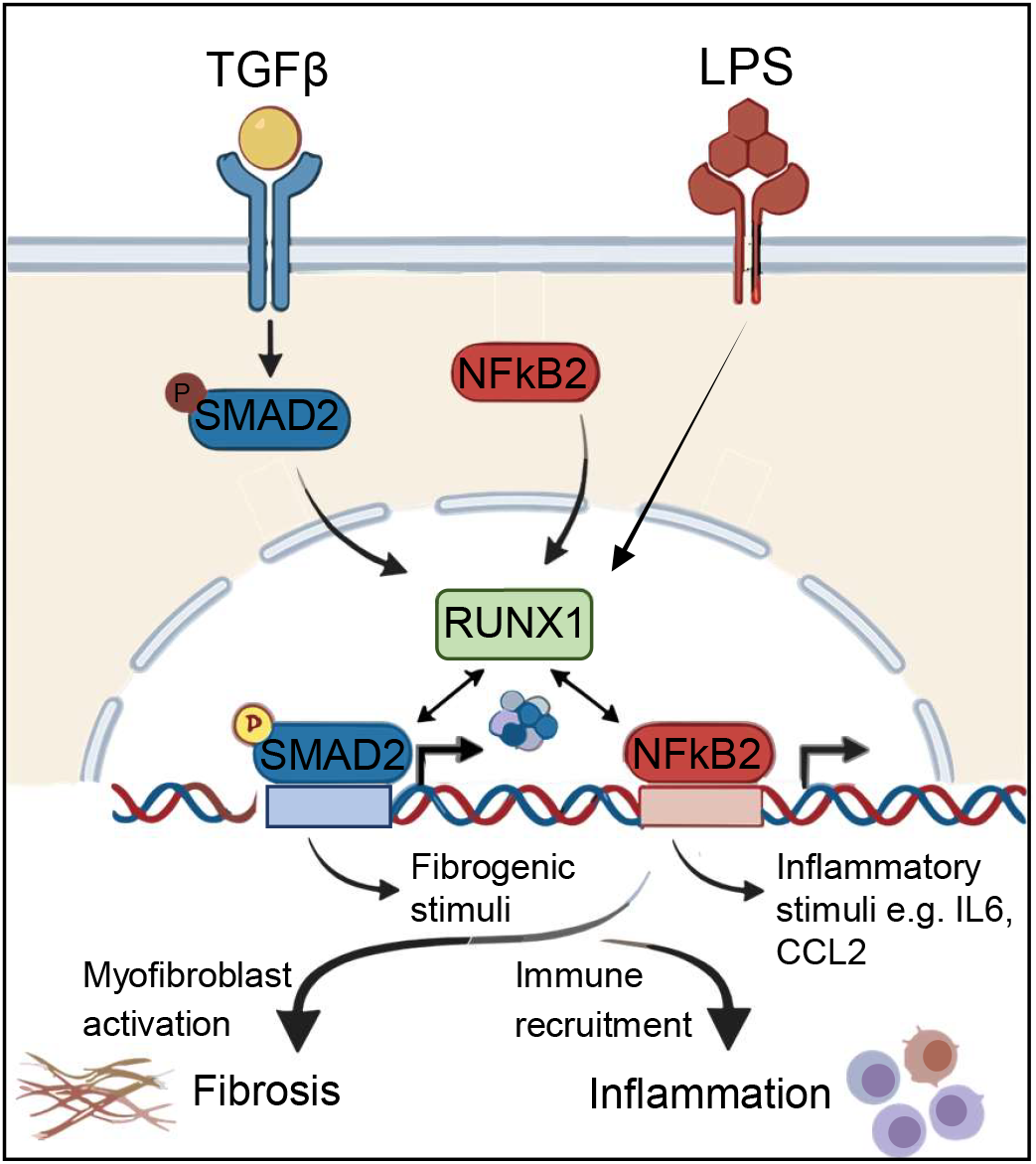
TGFβ activation upregulates RUNX1 in the nucleus, where it can interact with SMAD2 or NFKB2 signaling allowing for the expression of fibroinflammatory signals in cholangiocytes. RUNX1 is also essential in mediating the effects of inflammatory stimuli, such as LPS, in producing inflammatory signals.

## DISCUSSION

Biliary fibrosis is the predominant pathological driver in cholestatic liver diseases like PSC, yet therapeutic options remain limited [34, 35]. This is primarily due to an incomplete understanding of the pathogenesis of this process in these diseases. In this study, we identified the transcription factor RUNX1 as a novel and critical mediator of cholangiocyte-immune crosstalk during biliary inflammation and fibrosis. We demonstrated that RUNX1 is upregulated in human and murine cholestasis, driven by TGFβ as well as inflammatory signaling pathways, and is essential for the secretion of pro-inflammatory cytokines that propagate the fibroinflammatory response (Fig. 9).

In primary cholangiocytes derived from PSC patients, RUNX1 regulated the expression of prominently expressed IL6 and other inflammatory cytokines such as TNFα (Fig. 1). Similarly, RUNX1 regulated TGFβ-stimulated expression of IL6 by interacting with the canonical SMAD signaling in our cholangiocyte cell line. Indeed, RUNX1 directly bound the promoter regions of these genes. Bioinformatic analysis of these genes demonstrated the likelihood of additional upstream inflammatory regulators such as LPS, TNFα and NFKB, which were supported by assessing the role of RUNX1 in LPS- and TNFα-stimulated cholangiocytes. These findings suggest that RUNX1 is a promiscuous transcription factor hub, regulating and interacting with multiple fibrogenic and inflammatory pathways. Congruently, previous studies demonstrated RUNX1 mediating the downstream effects of TGFβ-SMAD signaling in HSCs and other tissues [18, 36, 37]. We also demonstrated that the TGFβ-RUNX1 pathway directly interacts with the non-canonical NF-κB2, which may be responsible for the expression of inflammatory mediators such as IL6 [38, 39]. It is possible that the TGFβ-RUNX1 signaling axis may involve additional pathways in upregulating fibroinflammatory signals. In contrast, LPS signaling has been shown to involve RUNX1 directly interacting with NFKB (p50) signaling [40, 41]. However, the outcome of this interaction is context and cell depending, for example activation of NFKB inflammatory signaling in macrophages while suppression of this pathway in neutrophils [40, 42]. In TNFα signaling, RUNX1 appears to mediate signaling via JNK-AP-1-RUNX1 axis [43].

While TGFβ is classically thought to be fibrogenic and anti-inflammatory, here we show that at least a component of TGFβ signaling is involved in perpetuating inflammatory mediators [44]. These observations are consistent with previous findings linking TGFβ with the NF-κB2 signaling in other epithelial cells [45]. However, the outcome of these interactions, i.e. pro- or anti-inflammatory effects, are highly context and disease dependent [44, 46, 47] [48]. TGFβ can attenuate the differentiation and activation of various immune components, while, depending on the signaling milieu, can promote differentiation and activation of immune cells, the prime example of which are the Th17 cells [46, 49]. Therefore, understanding the signaling pathways that are involved in this dual role of TGFβ will be valuable in therapeutic targeting. To that end, we show that RUNX1 is the critical co-mediator in the expression of TGFβ-stimulated fibroinflammatory stimuli in cholangiocytes, and biliary fibrosis and inflammation. RUNX1 interactions with the canonical NF-κB has been demonstrated in myeloid cells that can be either inhibiting or stimulating [40, 50]. However, RUNX1 interactions with the non-canonical NF-κB2 pathway was not previously documented. Taken together, these observations indicate that the TGFβ-RUNX1 axis interactions with the inflammatory canonical or non-canonical NF-κB signaling is highly context dependent and result in the stimulation of both fibrosis and inflammation in cholangiocytes in the context of biliary injury [2, 51].

Importantly, we show that the RUNX1 pathway is druggable. Treatment with the small molecule inhibitor Ro5-3335 effectively reduced fibrosis and inflammation in the Mdr2^−/−^ model, suggesting that repurposing RUNX1 inhibitors could be a viable therapeutic strategy for patients with biliary fibrosis. The anti-inflammatory effects of the RUNX1 inhibitor were prominent in the Mdr2^−/−^ mice on the white FVB/N background, which are known to develop more robust hepatitis [31-33]. This was evident in comparing the liver serum injury markers in both strains where the FVB/N strain developed significantly higher levels of serum AST/ALT that were significantly attenuated with Ro5-3335 (Fig. 6D). Although the end outcome was a broad suppression of inflammation in the liver, the effect of Ro5-3335 treatment was concentrated on CD4+ T cells, B cells and ITGAX+ group of myeloid cells. Since the effect of RUNX1 inhibition in the bile ducts cannot be distinguished from the direct effects on immune cells in a pharmacologic inhibition model, we also utilized a genetic model of specifically knocking down RUNX1 in cholangiocytes to demonstrate similar downregulation of inflammation and fibrosis in the DDC diet model. Our findings highlight a distinct role for RUNX1 in the biliary epithelium. While previous studies have implicated RUNX1 in hepatic stellate cell activation[18, 19, 36], our data using K19-Cre driven deletion shows that cholangiocyte-specific RUNX1 is required for the recruitment of inflammatory cells and the subsequent activation of myofibroblasts. This supports the “reactive cholangiocyte” hypothesis [52-54], where biliary epithelial cells are not just innocent bystanders but active drivers of disease progression through secretion of pro-inflammatory and pro-fibrogenic mediators[55, 56].

In contrast, RUNX1 knockdown using an albumin promoter-driven Cre resulted in worsened fibrosis and markers of inflammation in cholestatic mouse livers [20]. While these contradictory findings could be due to methodological differences, it is also possibly due to cell-specific roles of RUNX1 where in hepatocytes it might play a protective role but in cholangiocytes, HSCs and hepatic endothelial cells a paradoxical pro-fibroinflammatory one [18, 19, 36, 57, 58]. Furthermore, RUNX1 might also have important pro-fibroinflammatory roles in the immune compartment of biliary injury and fibrosis that has not been thoroughly examined [59]. Nonetheless, inhibition of RUNX1 in the liver as a whole using the small molecule inhibitor, Ro5-3335, attenuated biliary fibrosis and inflammation as we demonstrated in our Mdr2^−/−^ mouse strains, suggesting that the potential protective effects of RUNX1 are outweighed by its fibroinflammatory effects when inhibiting RUNX1 in the cholestatic liver. These findings are further supported by evidence from other solid organs, where RUNX1 also potentiates fibrosis [60].

In conclusion, this study demonstrated RUNX1 as a key orchestrator of the fibrogenic and inflammatory program in activated cholangiocytes. Targeting the TGFβ-RUNX1 axis offers a promising new avenue for treating PSC and other fibroinflammatory cholangiopathies.

## Acknowledgements

This study was funded in part by the **PSC Partners Seeking a Cure** Young Investigator Award (SOA) and Seed Grant (NJS) and NIH NIDDK **K08DK140618** Career Development Award (SOA). Additional funding came from the Stravitz-Sanyal Institute for Liver Disease and Metabolic Health, and the Department of Internal Medicine Pilot Fund, VCU. The data included in this study was generated at the Genomics Core facility at VCU. Services in support of the research project were provided by the VCU Massey Comprehensive Cancer Center Bioinformatics Shared Resource. Massey is supported, in part, with funding from NIH-NCI Cancer Center Support Grant P30 CA016059. HZ is supported by VA Merit Award I01BX005730, Research Career Scientist Award (IK6BX004477), VA ShEEP grants (1IS1BX005517-01 and 1 IS1 BX004777-01), National Institutes of Health Grant R01 DK104893, and R01DK-057543. We sincerely thank Dr. Daniel Goldenberg (Hadassah-Hebrew University Medical Center, Jerusalem, Israel) for providing the Mdr2^−/−^ mice and Dr. Nicholas LaRusso (Mayo Clinic, Rochester, MN, USA) for providing PSC-Cs.

## Figure legends

**Supplementary Table 1.** A) Mean RUNX1 expression level in RUNX1+ cholangiocytes as well as the fraction of RUNX1+ cholangiocytes in healthy, PSC and PBC livers (PSC/PBC sc/snRNAseq atlases and GEO database GSE243977). B) RUNX1 expression is significantly increased in RNA-seq analysis performed on liver samples from PSC subjects (GEO data set: GSE159676).

| Phenotype | Cell Type | Mean RUNX1+ | Percent Positive | Number of Cells |
| --- | --- | --- | --- | --- |
| Healthy | Cholangiocyte | 0.04 | 8.3 | 252 |
| PSC | Cholangiocyte | 0.12 | 47.2 | 362 |
| PBC | Cholangiocyte | 0.12 | 51.5 | 1067 |

B
| Gene | LogFc | p-value | FDR |
| --- | --- | --- | --- |
| RUNX1 | 15.171 | 0.002 | 0.015 |
| RUNX1T1 | 0.123 | 0.732 | 1.000 |
| RUNX2 | 0.168 | 0.614 | 1.000 |
| RUNX3 | 0.037 | 0.907 | 1.000 |

**Supplementary Table 2.**
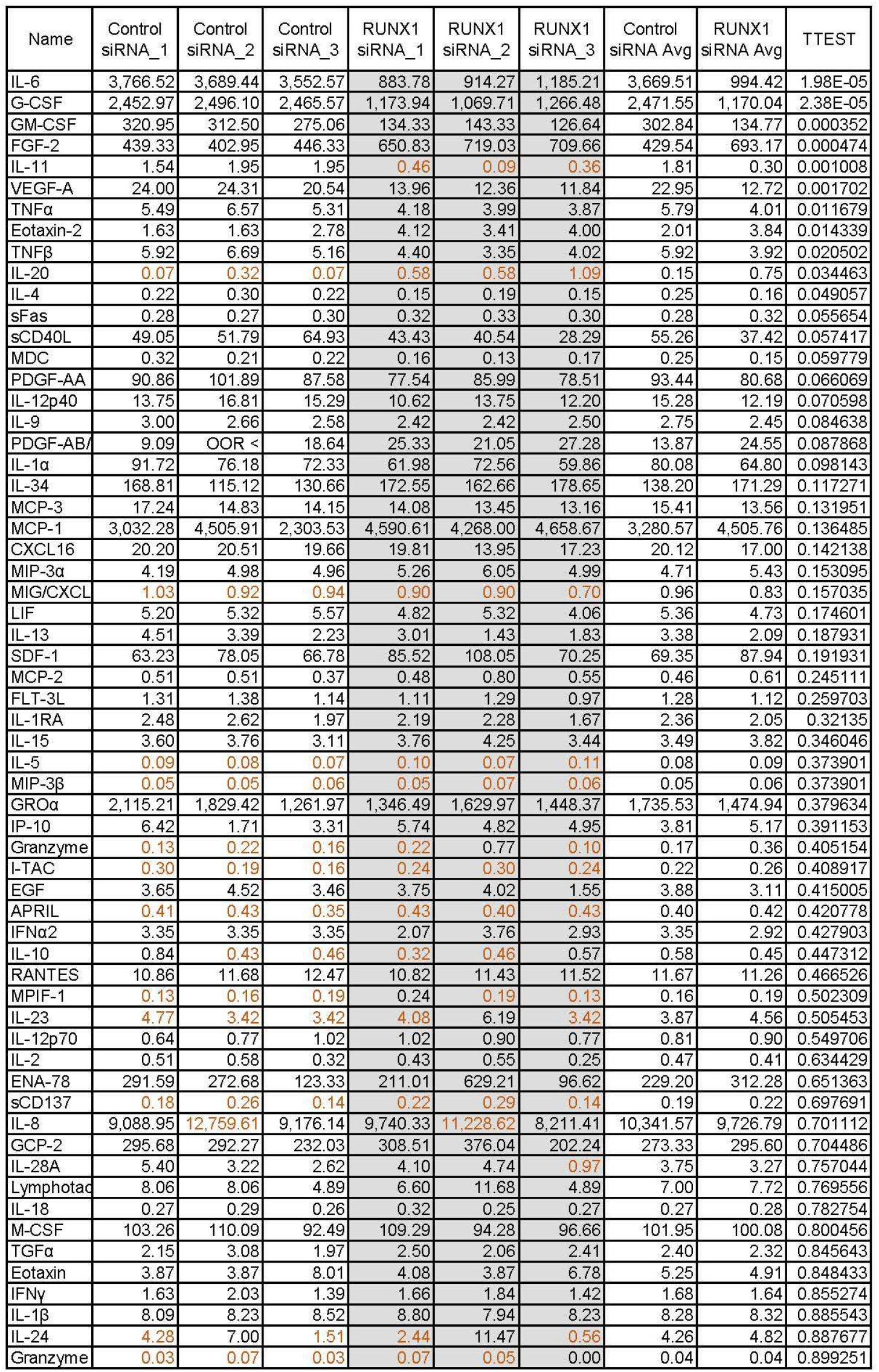
Levels of cytokines in the media of PSC-Cs treated with control or RUNX1 siRNA measured using Luminex xMAP technology based multiplexed quantification (pg/mL).

**Supplementary Fig 1.**
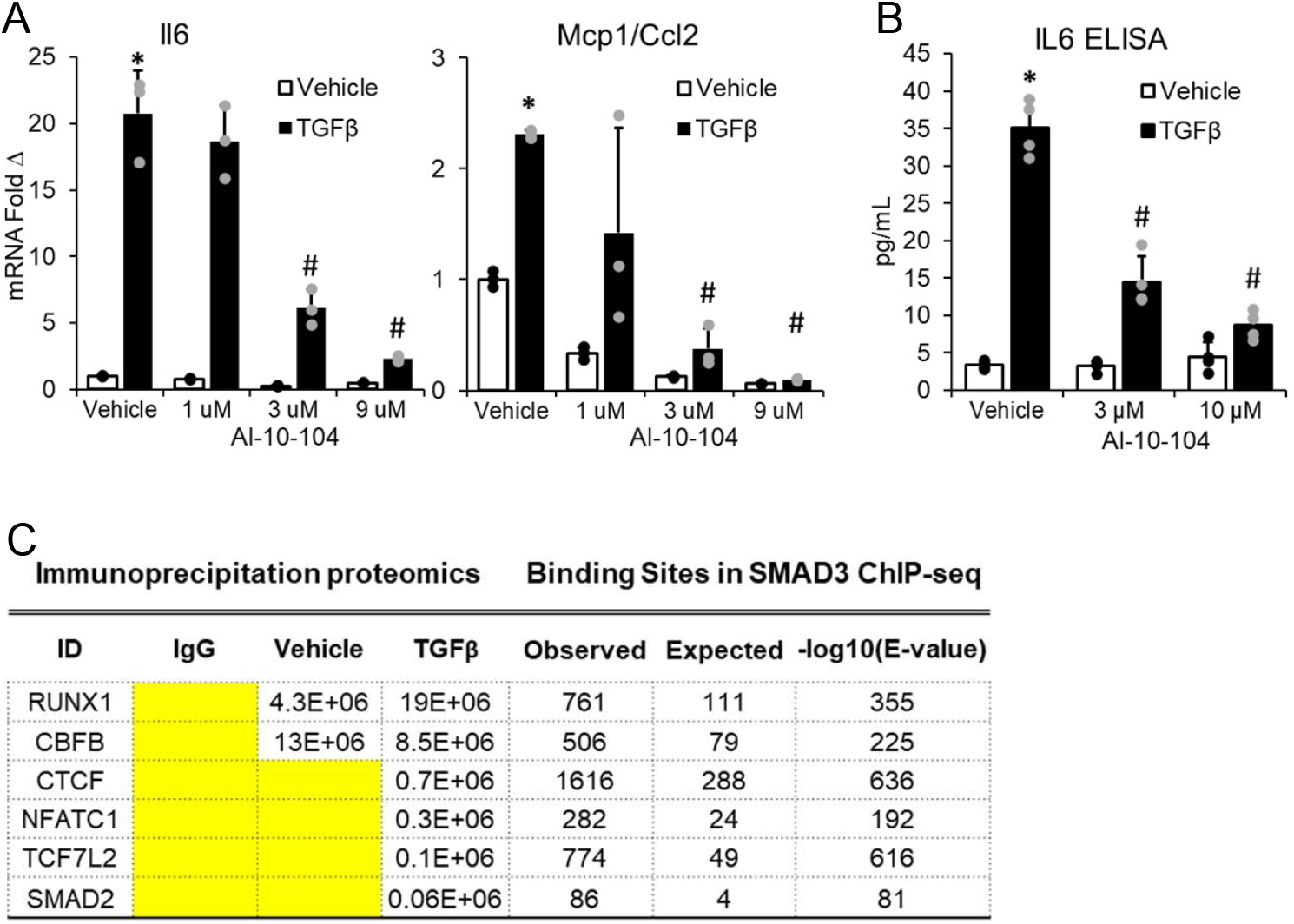
In MLEs, the TGFβ-induced elevated expression of *Il6* and *Ccl2/Mcp1* mRNA expression and IL6 protein level were dose-dependently attenuated with the RUNX1 inhibitor, AI-10-104, co-treatment (A-B). (N=3 at least, # p<0.05 when compared to TGFβ only)

**Supplementary Fig 2.**
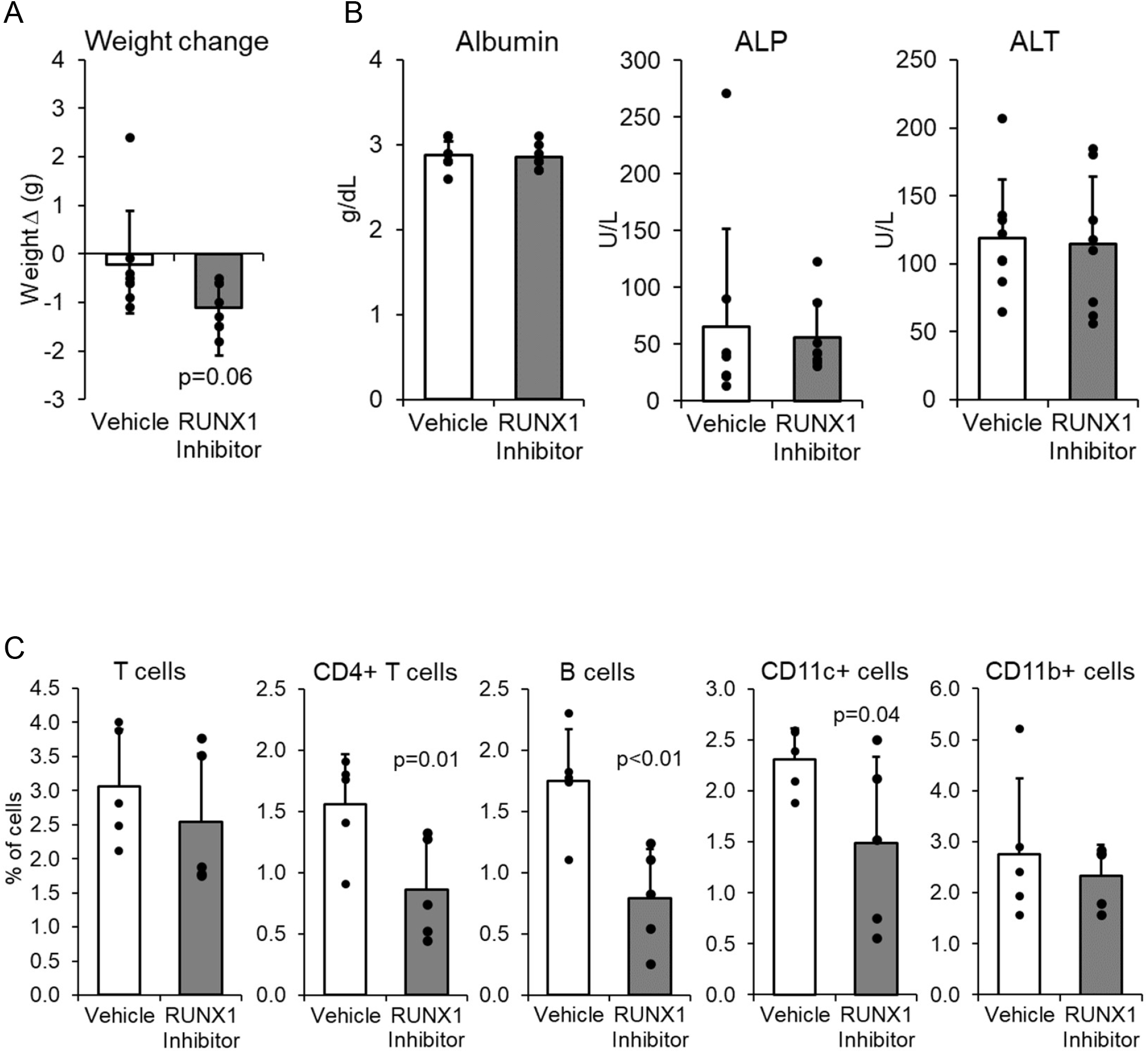
A) Mdr2^−/−^ mice on the C57/B6 background treated with Ro5-3335 appeared to have more weight loss approaching statistical significance compared to vehicle treated mice. B) Serum hepatic injury markers, alkaline phosphatase (ALP), alanine aminotransferase (ALT), or serum albumin were not significantly different between Ro5-3335 and vehicle only treated mice. (N= 8 per group). C) In Mdr2^−/−^ mice on the FVB/N background, fraction of immune cells in the peri-portal regions determined by IF staining for the markers indicated. (N= 5 per group)

**Supplementary Fig 3.**
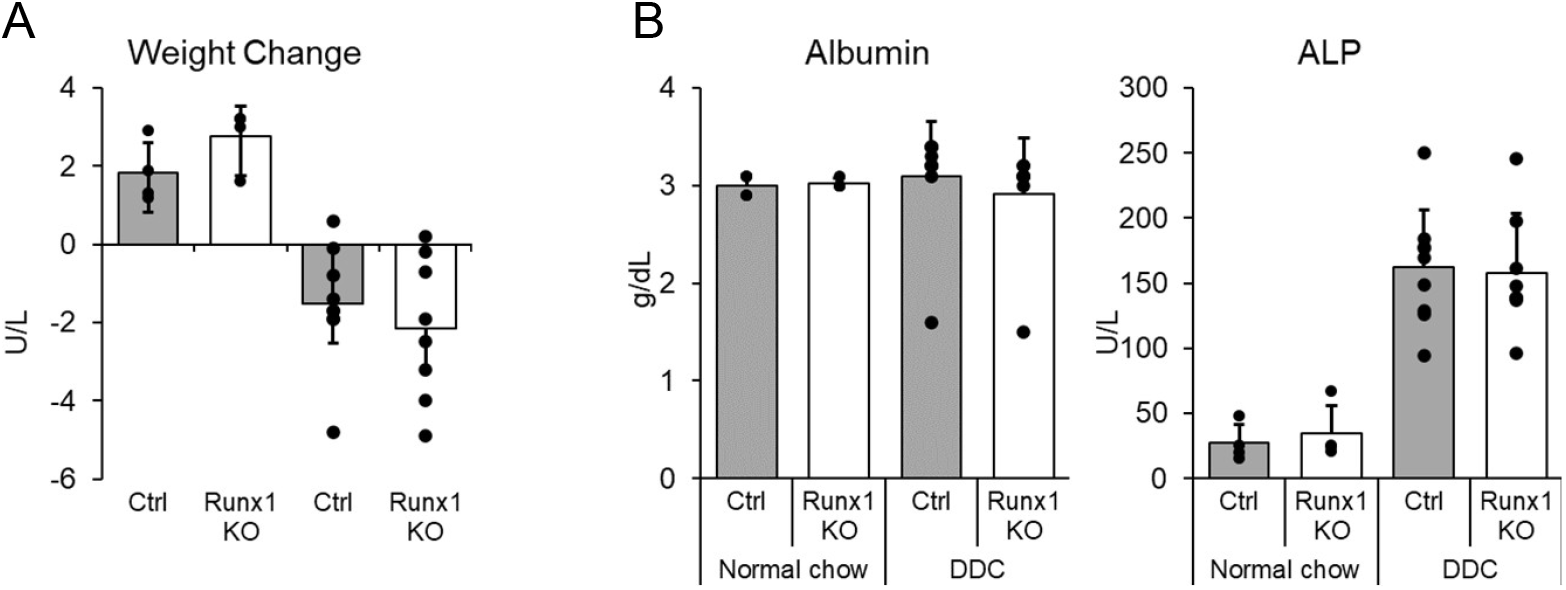
A) DDC diet induced weight changes were not significantly different in cholangiocyte-specific *Runx1* deletion (*Runx1* KO) compared to control mice. B) Serum albumin and alkaline phosphatase (ALP) were not significantly different in *Runx1* KO compared to control mice. (N=4 per regular chow group, 8 per DDC diet group).

